# Whole Root Transcriptomic Analysis Reveals a Role for Auxin Pathways in Resistance to *Ralstonia solanacearum* in Tomato

**DOI:** 10.1101/176685

**Authors:** Elizabeth French, Bong Suk Kim, Katherine Rivera-Zuluaga, Anjali S. Iyer-Pascuzzi

## Abstract

The soilborne pathogen *Ralstonia solanacearum* is the causal agent of bacterial wilt, and causes significant crop loss in the Solanaceae family. The pathogen first infects roots, which are a critical source of resistance in tomato (*Solanum lycopersicum* L.). Roots of both resistant and susceptible plants are colonized by the pathogen, yet rootstocks can provide significant levels of resistance. Currently, mechanisms of this ‘root-mediated resistance’ remain largely unknown. To identify the molecular basis of this resistance, we analyzed the genome-wide transcriptional response of roots of resistant (Hawaii 7996) and susceptible (West Virginia700) tomatoes at multiple time points after inoculation with *R. solanacearum*. We found that defense pathways in roots of the resistant Hawaii7996 are activated earlier and more strongly than roots of susceptible West Virginia700. Further, auxin signaling and transport pathways are suppressed in roots of the resistant variety. Functional analysis of an auxin transport mutant in tomato confirmed a role for auxin pathways in bacterial wilt. Together, our results suggest that roots mediate resistance to *R. solanacearum* through genome-wide transcriptomic changes that result in strong activation of defense genes and alteration of auxin pathways.

## INTRODUCTION

The soilborne betaproteobacterium *Ralstonia solanacearum* is the causal agent of bacterial wilt and has been ranked as one of the top 10 most destructive plant bacterial pathogens of all time (Mansfield et al. 2012). The pathogen infects over 200 plant species in 50 families, but is particularly devastating to members of the Solanaceae family (Hayward 1991; Huet 2014). *R. solanacearum* is a vascular pathogen that first colonizes the root surface and subsequently enters the root of both resistant and susceptible plants through small natural wounds or root tips (Genin 2010). The bacterium secretes cell wall-degrading enzymes and eventually spreads into the vascular system where it moves to the shoot via the flow of xylem fluid (Genin 2010; Genin and Denny 2012). As bacteria multiply, they secrete exopolysaccharide (EPS) (Genin 2010; Genin and Denny 2012), which likely leads to physical xylem blockage, and aboveground wilting. Resistant plants are able to delay colonization of the root vasculature (Caldwell et al. 2017), but the molecular responses involved in this delay are not clear. Here we use RNA-seq and mutant analysis to understand responses to *R. solanacearum* in roots of resistant and susceptible tomato genotypes.

In tomato, resistance to *R. solanacearum* is quantitative (Danesh et al. 1994; Thoquet et al. 1996a; Thoquet et al. 1996b; Wang et al. 2000; Carmeille et al. 2006; Wang et al. 2013; Kim et al. 2016), but no quantitative trait loci (QTL) for resistance have been cloned. Microarray analysis of genes differentially expressed in tomato stems 24 hours after infection showed that *R. solanacearum* activates defense, hormone, and lignin pathways in resistant tomato stems (Ishihara et al. 2012). Surprisingly, no differentially expressed genes (fold change > 2) were identified in susceptible stems after infection (Ishihara et al. 2012).

Despite the prevalence of soilborne pathogens and root diseases, most work in plant-pathogen interactions has focused on the aboveground portion of the plant. This is likely due to the hidden nature of roots, and the visible aboveground disease phenotypes that often result from root infection. However, recent reports indicate that roots also have a robust immune system that functions to protect the plant from soilborne pathogens. For example, Arabidopsis roots can recognize microbe associated molecular patterns (MAMPs) from pathogenic bacteria (Millet et al. 2010). In addition, roots infected with nematodes, which colonize root cortex tissue, can activate both MAMP-triggered immunity (MTI) (Teixeira et al. 2016) and effector-triggered immunity (ETI) (Mitchum et al. 2013; Goverse and Smant 2014). Tomato roots also appear to mount a defense response to *R. solanacearum* because resistant rootstocks grafted to susceptible scions result in scions that are resistant to *R. solanacearum* and do not wilt (McAvoy et al. 2012; Rivard et al. 2012).

One approach to uncover the mechanisms of resistance in tomato roots to *R. solanacearum* is the analysis of whole genome transcriptional responses. In resistant and susceptible accessions of a wild potato species, *Solanum commersonii*, transcriptome analysis 3 – 4 days after inoculation with *R. solanacearum* identified 221 genes in the resistant accession and 644 genes in the susceptible that respond to infection (Chen et al. 2014; Zuluaga et al. 2015). In both accessions, genes that function in development were primarily downregulated, while those in the gene ontology category ‘biotic stress’ were mainly upregulated after infection (Zuluaga et al. 2015). In contrast, in a timecourse of peanut root infection the expression patterns of many defense genes, including LRR-Kinases and *R* genes were mainly downregulated in both resistant and susceptible peanut genotypes (Chen et al. 2014). Carbohydrate metabolism was repressed after infection in roots of both resistant and susceptible peanut roots, but more strongly inhibited in resistant roots (Chen et al. 2014). This suggests that the mechanisms of root-mediated resistance may differ among plant species.

The plant hormone auxin can have both positive and negative effects on plant defense, (reviewed in (Kazan and Manners 2009; Fu and Wang 2011; Ludwig-Muller 2015)). Plant resistance to some necrotrophic pathogens requires auxin signaling (Tiryaki and Staswick 2002; Llorente et al. 2008; Qi et al. 2012), but multiple reports have revealed a relationship between plant susceptibility to biotrophic pathogens and increased auxin accumulation or signaling (O’Donnell et al. 2003; Navarro et al. 2006; Chen et al. 2007; Ding et al. 2008; Fu and Wang 2011). Many phytopathogens produce auxin (Spaepen et al. 2007; Ludwig-Muller 2015), and this probably includes *R. solanacearum* (Valls et al. 2006). Exogenous treatment with auxin or auxin analogs increases disease symptoms caused by *Pseudomonas syringae* in Arabidopsis (Navarro et al. 2006; Chen et al. 2007) and increases rice susceptibility to *Xanthomonas oryzae* pv. *oryzae* (Ding et al. 2008), *X. oryzae* pv *oryzicola* and *Magnaporthe grisea* in rice (Fu and Wang 2011). In Arabidopsis, overexpression of the AvrRpt2 type III effector from *Pseudomonas syringae* changes auxin-related developmental phenotypes (Chen et al. 2007) through the ability of AvrRpt2 to promote degradation of an AUX/IAA transcription factor, AXR2/IAA7, which represses auxin responses (Cui et al. 2013).

Suppression of auxin signaling may be particularly important in plant defense against vascular wilt pathogens. Several Arabidopsis auxin signaling and transport mutants are resistant to the soilborne vascular wilt pathogen *Fusarium oxysporum* (Kidd et al. 2011), and the walls are thin (*wat1*) mutant of Arabidopsis is resistant to multiple vascular wilt pathogens, including *R. solanacearum* (Denance et al. 2012). The *wat1* mutant has decreased levels of auxin in roots (Denance et al. 2012) and the base of stems (Ranocha et al. 2013), and the gene was recently shown to encode a vacuolar auxin transporter. *WAT1* is expressed in the root pericycle and lateral root primordium (Denance et al. 2012), suggesting that auxin homeostasis within these tissues is particularly important for bacterial wilt resistance.

In this study, we aimed to identify the transcriptional response of resistant and susceptible tomato roots to *R. solanacearum* infection at 24 hours and 48 hours post inoculation (hpi). We identified the responsive genes in resistant and susceptible accessions independently and compared the responses. We show that resistant tomato roots activate defense pathways and terpene biosynthesis genes, and suppress auxin signaling and transport pathways in response to *R. solanacearum*. In contrast, susceptible tomato roots activate defense response marker genes later, and at a lower fold change, and genes required for root growth are suppressed by 48 hours post inoculation. Consistent with our finding that auxin pathways are suppressed in resistant roots, we show that an auxin transport mutant in a susceptible tomato wild-type background is resistant to *R. solanacearum*. Our data suggest that tomato roots mediate resistance to *R. solanacearum* in part through the suppression of auxin pathways.

## RESULTS

### Roots of resistant and susceptible tomato plants have a strong transcriptional response to R. solanacearum infection

We utilized resistant Hawaii 7996 and susceptible West Virginia (WV) for our analyses. H7996 is a variety of cultivated tomato (*S. lycopersicum*) that is resistant to many different *R. solanacearum* strains (Lebeau et al. 2011). WV is an accession of *S. pimpinellifolium*, the closest wild relative to *S. lycopersicum* (Tomato Genome Sequencing Consortium 2012), and is highly susceptible to *R. solanacearum*. We chose these genotypes for transcriptomic analysis because they are the parents of a recombination inbred line population that has been used in multiple QTL (quantitative trait loci) studies (Thoquet et al. 1996a; Wang et al. 2000; Carmeille et al. 2006; Wang et al. 2013) for resistance to *R. solanacearum*. Transcriptomic data may be useful towards the further identification of genes underlying resistance QTL. Because resistant H7996 (*S. lycopersicum*) and susceptible WV (*S. pimpinellifolium*) are different species, we identified the response within each species by comparing each time point (24 hpi or 48 hpi) to the 0 hpi mock control for each genotype.

We hypothesized that transcriptional events that promoted defense responses in roots of resistant plants would occur early, before wilting, but would be non-existent or diminished in roots of susceptible plants. We inoculated roots using our previously established soil-soak inoculation method (Caldwell et al. 2017), in which wilting typically begins at 72 – 96 hpi in WV. We previously performed light and scanning electron microscopy and showed that bacteria colonize the root of both resistant H7996 and susceptible WV at 24 hpi and 48 hpi at 2.5 cm below the root-shoot junction (Caldwell et al. 2017). Here, we first tested whole roots to confirm that *R. solanacearum* colonizes roots of resistant H7996 and susceptible WV at 24 and 48 hpi (Fig. 1). Plants were grown in potting mix and inoculated with 10^8^ CFU/ml *R. solanacearum* K60 at the three-leaf stage as in (Caldwell et al. 2017). Consistent with our previous results, in three independent experiments, bacteria colonized roots of both resistant H7996 and susceptible WV at 24 and 48 hpi (Fig. 1). We then used genome wide-RNA seq analysis to identify the *R. solanacearum*-responsive transcriptome of whole roots in resistant H7996 and susceptible WV tomatoes prior to the onset of wilting at 0, 24, and 48 hpi. Plants were grown and root inoculated as above. Whole roots were harvested at 0, 24 and 48 hpi. Total RNA from 10 roots was pooled for each genotype at each time point and was sent to the Purdue Genomics Facility for library creation and sequencing on the Illumina HiSeq2500 (see Materials and Methods). Reads were mapped by the Purdue Genomics Facility using TopHat2 version 2.0.14 to the *S. lycopersicum* genome (ITAG2.4).

**Fig. 1:**
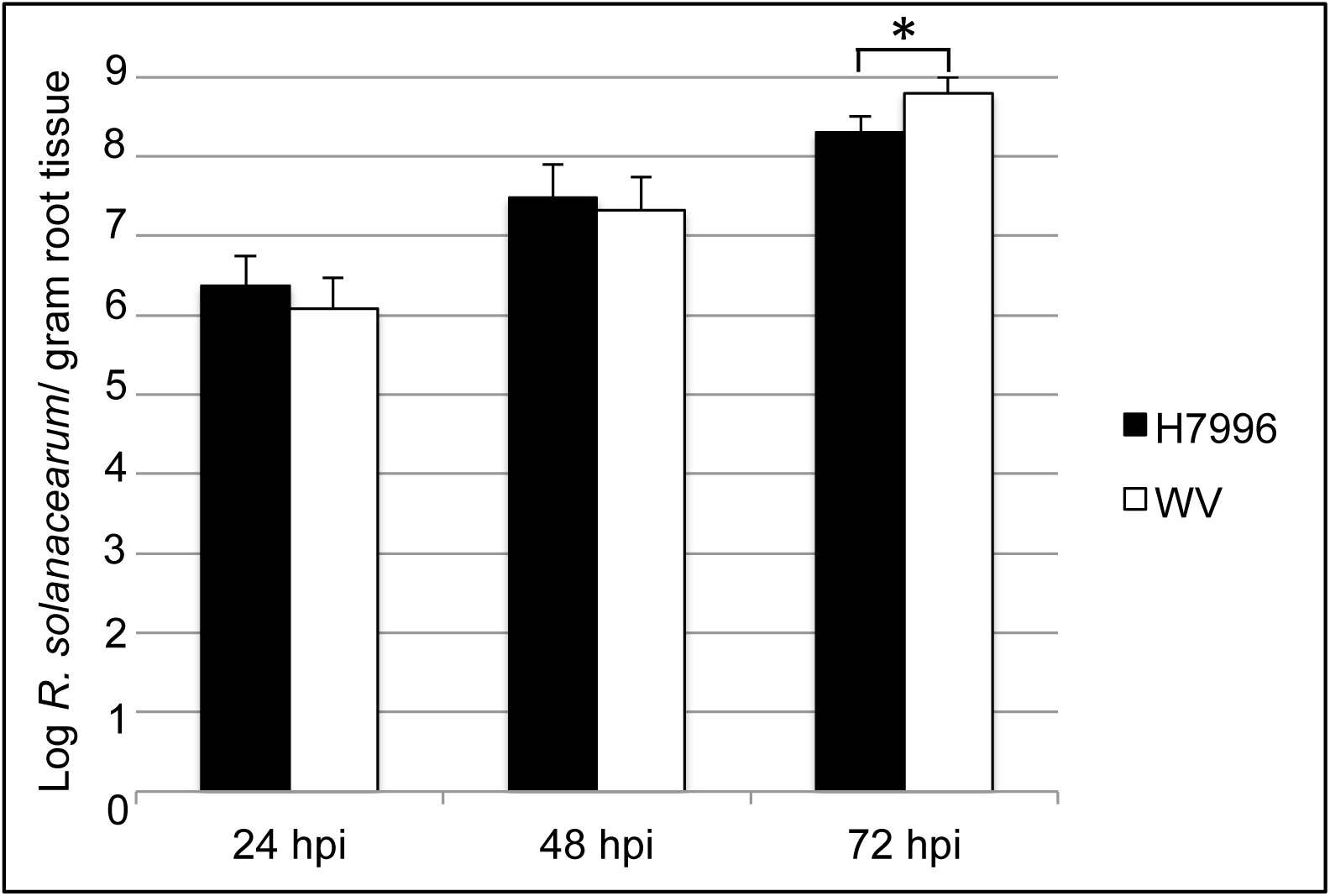
Root colonization of *R. solanacearum* K60 in whole roots of resistant H7996 and susceptible WV. Plants were grown in potting mix and root inoculated via soil soaking at the three-leaf stage. The average of three independent replicates, each with roots of three plants per genotype and time point, is shown. Error bars indicate standard deviation. * = P < 0.05 with the Mann Whitney Wilcoxon test.

Pairwise comparisons were made between each time point and 0 hpi (mock inoculated control) to identify transcriptional responses to *R. solanacearum* infection within each genotype. We classified responsive genes (hereafter ‘differentially expressed genes’ or DEGs) as those that showed a log2 fold change > |0.585| and a false discovery rate (FDR) < 0.05. To understand how the response to *R. solanacearum* infection in resistant and susceptible roots differed, the DEGs at each time point within a genotype were then compared between genotypes (Fig. 2). The mapping summary is in Supplementary Table S1, normalized library sizes are in Supplementary Table S2, raw counts are listed in Supplementary Table S3, and processed edgeR gene expression results are in Supplementary Table S4. Differential expression analysis showed that within susceptible roots at 24 hpi, 427 genes were upregulated and 545 downregulated, while within resistant roots at that time point almost twice as many genes were differentially expressed (957 up and 1029 down) (Fig. 2). At 48 hpi, 1316 genes were upregulated in susceptible roots and 1571 were downregulated compared to 1265 upregulated in resistant roots and 1419 downregulated. We used quantitative RT-PCR to validate the differential expression of fifteen genes. These showed similar expression patterns as identified in our RNA-seq analysis (Supplementary Fig. S1).

**Fig. 2:**
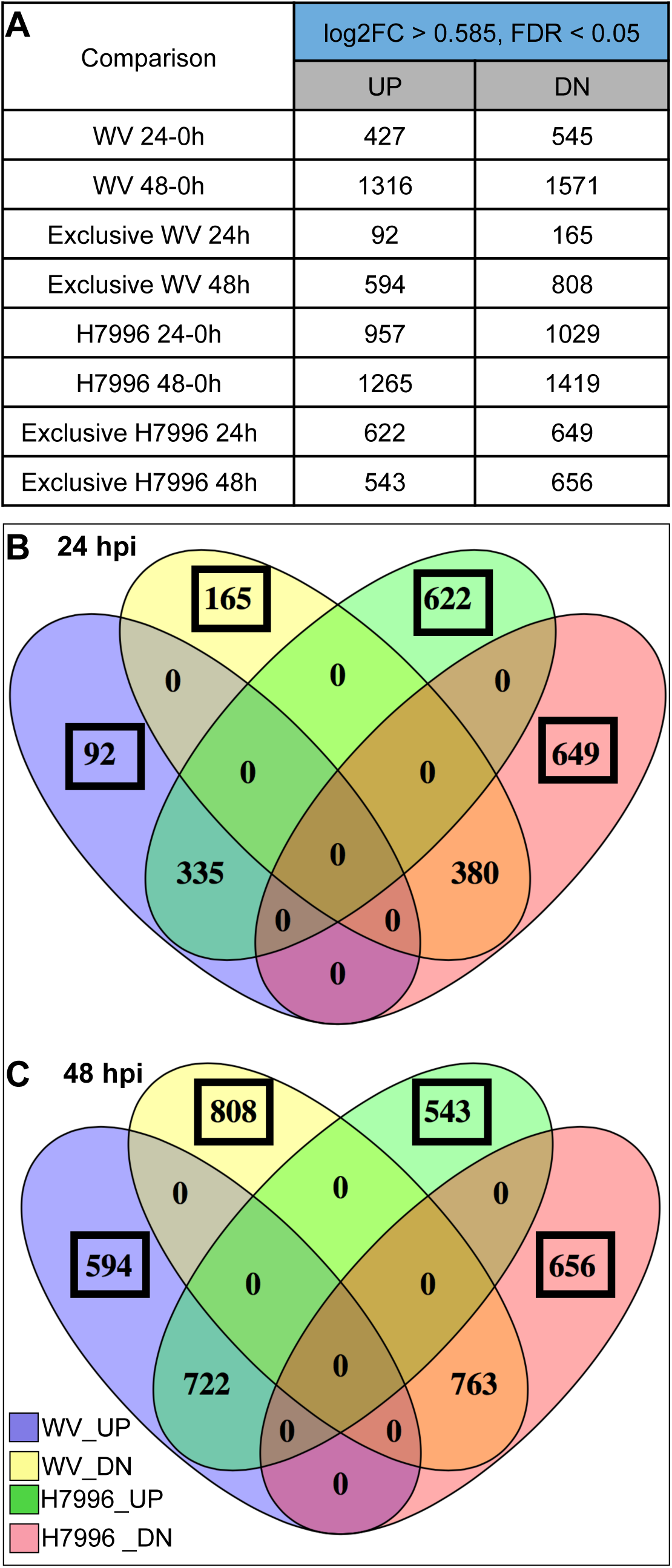
Summary of DEGs from pairwise comparisons between time points within each genotype (H7996 or WV). A) Numbers of DEGs at each pairwise comparison within each genotype. Threshold for differential expression is log2 fold change > |0.585|, False Discovery Rate (FDR) < 0.05. (B and C) Venn Diagram of up- and downregulated DEGs at 24 hpi (B) and 48 hpi (C) showing overlap between the responses of resistant *Solanum lycopersicum* L.) variety Hawaii7996 (H7996) and susceptible *S. pimpinellifolium* West Virginia 700 (WV). Boxed numbers show ‘exclusive’ genes at each time point.

At each time point, we also examined genes that were up or downregulated only within resistant H7996 or susceptible WV roots (Fig. 2b, 2c, boxed numbers). We call these genes ‘exclusive’ genes. Major shifts in numbers of exclusive DEGs were observed in susceptible roots between 24 and 48 hpi. For example, at 24 hpi, only 92 genes were exclusively upregulated in susceptible WV roots, compared to 622 genes in resistant H7996 roots. However, by 48 hpi, this number rose to 594 genes in susceptible WV roots compared to 543 in resistant H7996 roots (Fig. 2). We did not identify any significant DEGs whose expression was upregulated in roots of resistant H7996 and simultaneously downregulated in susceptible WV (or vice versa) at either time point.

We used Gene Ontology (GO) analysis to understand what biological processes were affected within roots of resistant H7996 and susceptible WV plants after inoculation. GO analysis using PANTHER (Huaiyu et al. 2016) showed that in susceptible WV at 24 hpi, only seven GO terms for biological process are overrepresented (P < 0.05) among the 427 genes upregulated (Supplementary Table S5). These include ‘response to stress’ (GO:0006950; P = 9.76 × 10^-3^) and ‘response to stimulus’ (GO:0050896; P = 2 × 10^-2^). In contrast, at 24 hpi in roots of the resistant H7996, 27 biological process GO terms were overrepresented in the 957 upregulated genes (Fig. 3 shows a subset of overrepresented GO categories, all overrepresented GO categories for all comparisons are in Supplementary Table S5). These included ‘reactive oxygen species metabolic process’ (GO:0072593; P = 6.3 × 10^-6^) and ‘cellular detoxification’ (GO:1990748; P = 8.7 × 10^-6^). Not unexpectedly, the GO category ‘defense responses’ (GO: 0006952; P = 2.45 × 10^-5^) was identified in upregulated genes in roots of the resistant plant at 24 hpi (Fig. 3), but was not present in upregulated genes of susceptible roots at this time point.

**Fig. 3:**
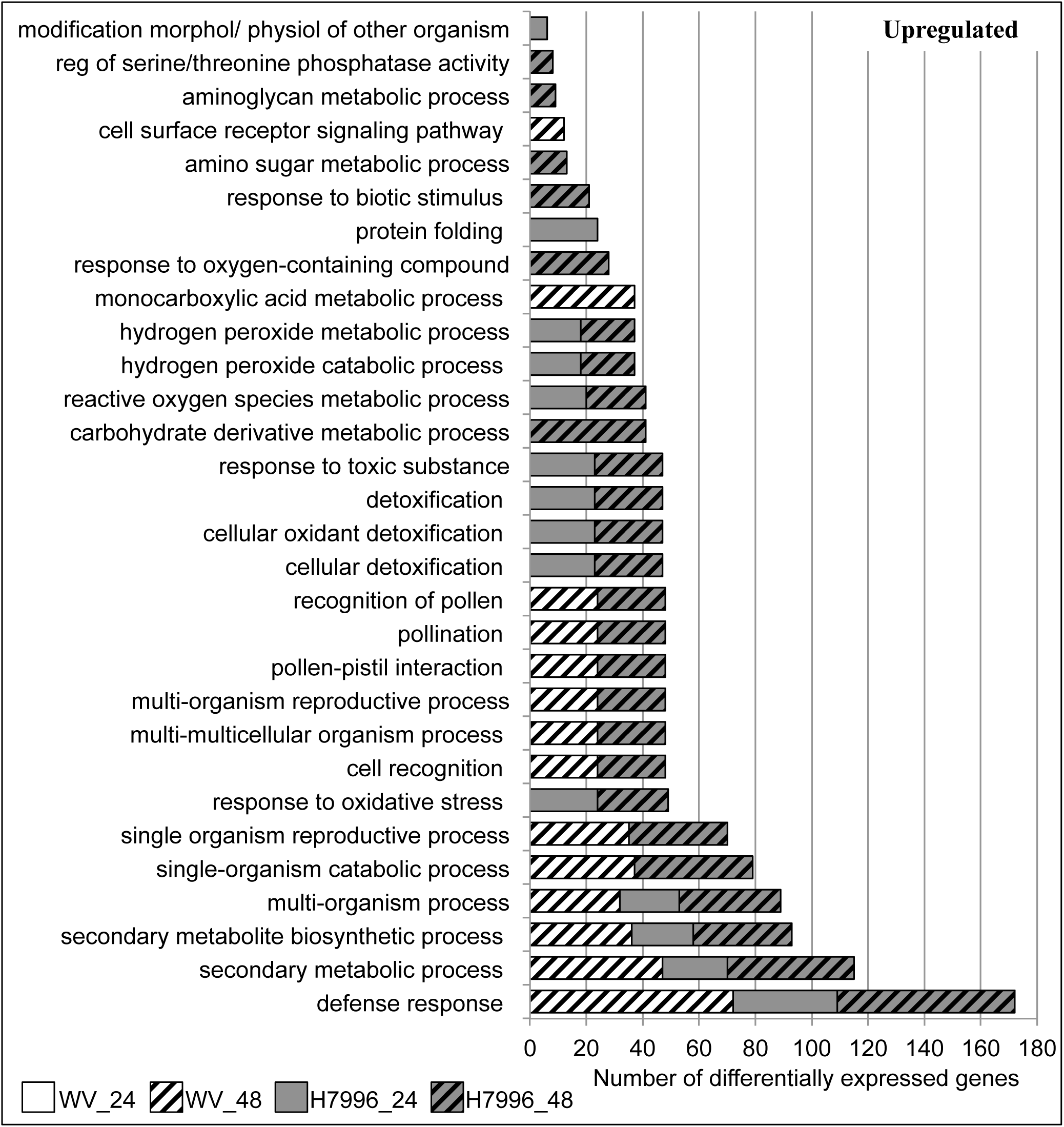
GO categories overrepresented (corrected P-value < 0.05) in the set of upregulated genes at each time point. Only categories that contain less than 600 total *S. lycopersicum* genes are shown in the figure (all categories are in Supplementary Table S5). WV 24 = 24 – 0 hpi comparison, WV 48 = 48 – 0 hpi comparison etc. No GO categories with less than 600 total genes are overrepresented in WV_24 upregulated genes.

Twenty-five biological process GO terms are overrepresented in the 545 downregulated genes at 24 hpi in susceptible WV roots, including ‘plant-type cell wall organization or biogenesis’ (GO:0071669; P = 2.38 × 10^-2^), ‘reactive oxygen species metabolic process’ (GO:0072593; P = 3.34 × 10^-3^), and ‘cellular detoxification’ (GO:1990748; P = 1.69 × 10^-4^) (Fig. 4 and Supplementary Table S5). Notably, and as stated above, the latter two GO categories were both overrepresented in *upregulated* genes in resistant roots at this time point. GO overrepresentation in downregulated H7996 genes at 24 hpi included ‘regulation of jasmonic acid (JA) mediated signaling pathway’ (GO:2000022; P = 1.26 × 10^-6^) (Fig. 4), consistent with the downregulation of JA responses in resistant plants after infection with some biotrophic pathogens (Spoel et al. 2003; Glazebrook 2005; Spoel et al. 2007; Koornneef et al. 2008; Koornneef and Pieterse 2008).

**Fig. 4:**
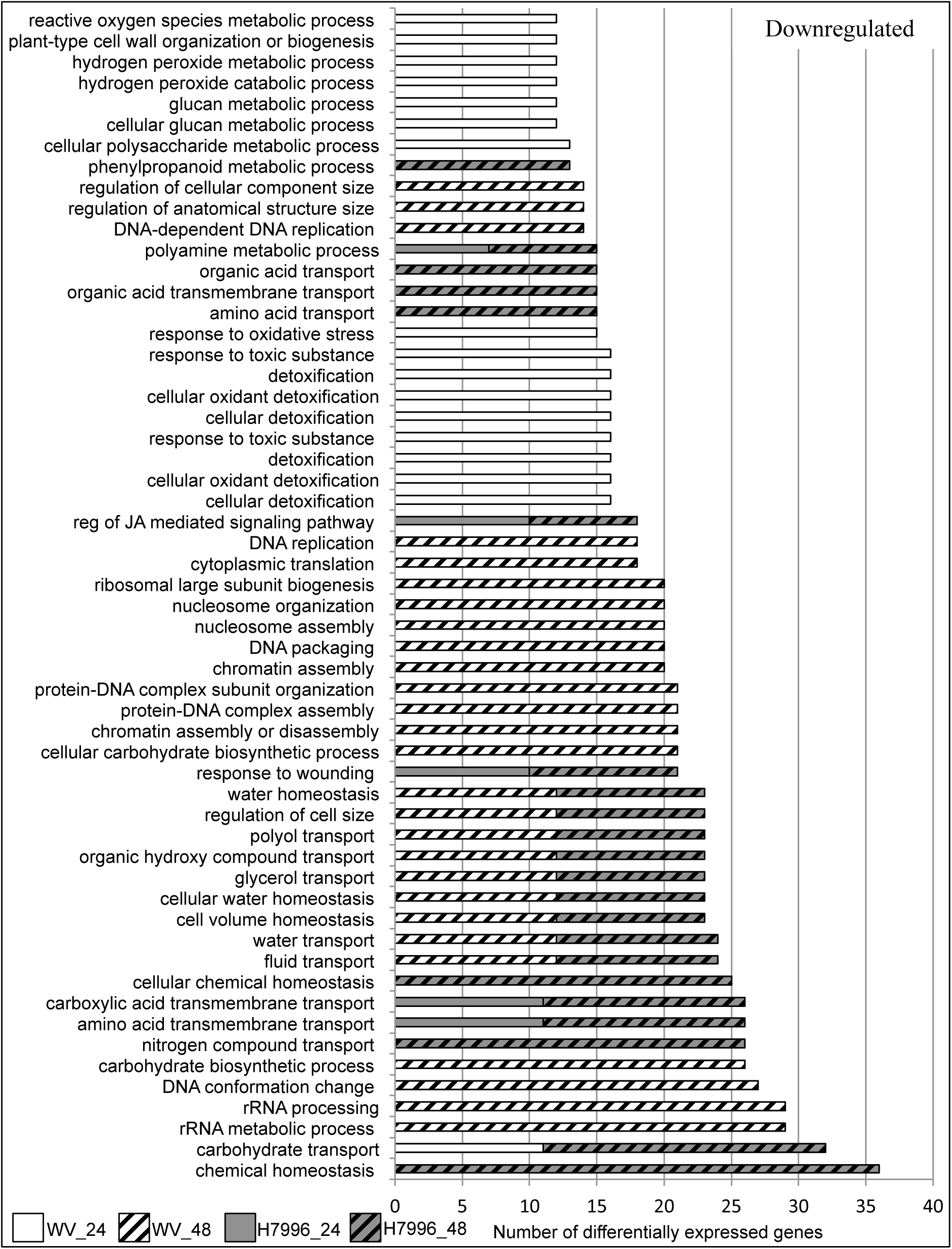
GO categories overrepresented (corrected P-value < 0.05) in the set of downregulated genes at each time point. Only categories that contain less than 300 total *S. lycopersicum* genes are shown in the figure (all categories are in Supplementary Table S5). WV 24 = 24 – 0 hpi comparison, WV 48 = 48 – 0 hpi comparison etc.

Many of the same trends in GO terms were observed at 48 hpi as at 24 hpi in each genotype. For example, ‘Reactive oxygen species metabolic process’ and ‘cellular detoxification’ categories were still overrepresented in upregulated genes in the resistant H7996 root at 48 hpi (Fig. 3) (P = 5.41 × 10^-5^ and P = 3.47 × 10^-4^, respectively), but were not overrepresented in upregulated genes of the susceptible WV root at either time point (Fig. 3). The GO category ‘defense response’ continued to be overrepresented in upregulated genes of the resistant H7996 root at 48 hpi (P = 2.98 × 10^-15^) (Fig. 3). While the ‘defense response’ category was not overrepresented at 24 hpi in the root of susceptible WV, it was identified at 48 hpi (P = 4.27 × 10^-20^) in upregulated genes of the susceptible WV root (Fig. 3). In downregulated genes, ‘Cell wall organization or biogenesis (GO:0071554)’ was overrepresented in susceptible roots at 48 hpi (P = 1.46 × 10^-4^) (see Supplementary Table S5), while ‘JA mediated signaling pathway’ continued to be overrepresented in the resistant H7996 plant at 48 hpi (P = 3.78 × 10^-3^) (Fig. 4).

### Defense gene activation occurs earlier and is stronger in roots of resistant tomato plants

Our GO analysis of genes up and downregulated at each time point showed that roots of resistant plants activated genes enriched for immune GO categories (such as ‘response to biotic stimulus’, response to oxidative stress’, ‘defense response’, and response to stimulus) earlier in the resistant H7996 root than in the susceptible WV root (Fig. 3 and 4).

To examine this more carefully, we next focused on the expression of specific defense marker genes in classic defense hormone pathways. We examined genes previously used as markers for defense responses in resistant H7996 (Milling et al. 2011). The ethylene (ET) marker gene *PR-1b* was upregulated only in the resistant H7996 genotype, while *Osmotin* was activated earlier and with a higher fold change compared to 0 hpi in H7996 compared to WV (Fig. 5a). SA marker genes were similarly regulated, with *PR-1a* being exclusively activated in H7996 at 48 hpi, and *Glu-A* was activated more strongly in H7996 compared to susceptible WV at both 24 and 48 hpi.

**Fig. 5:**
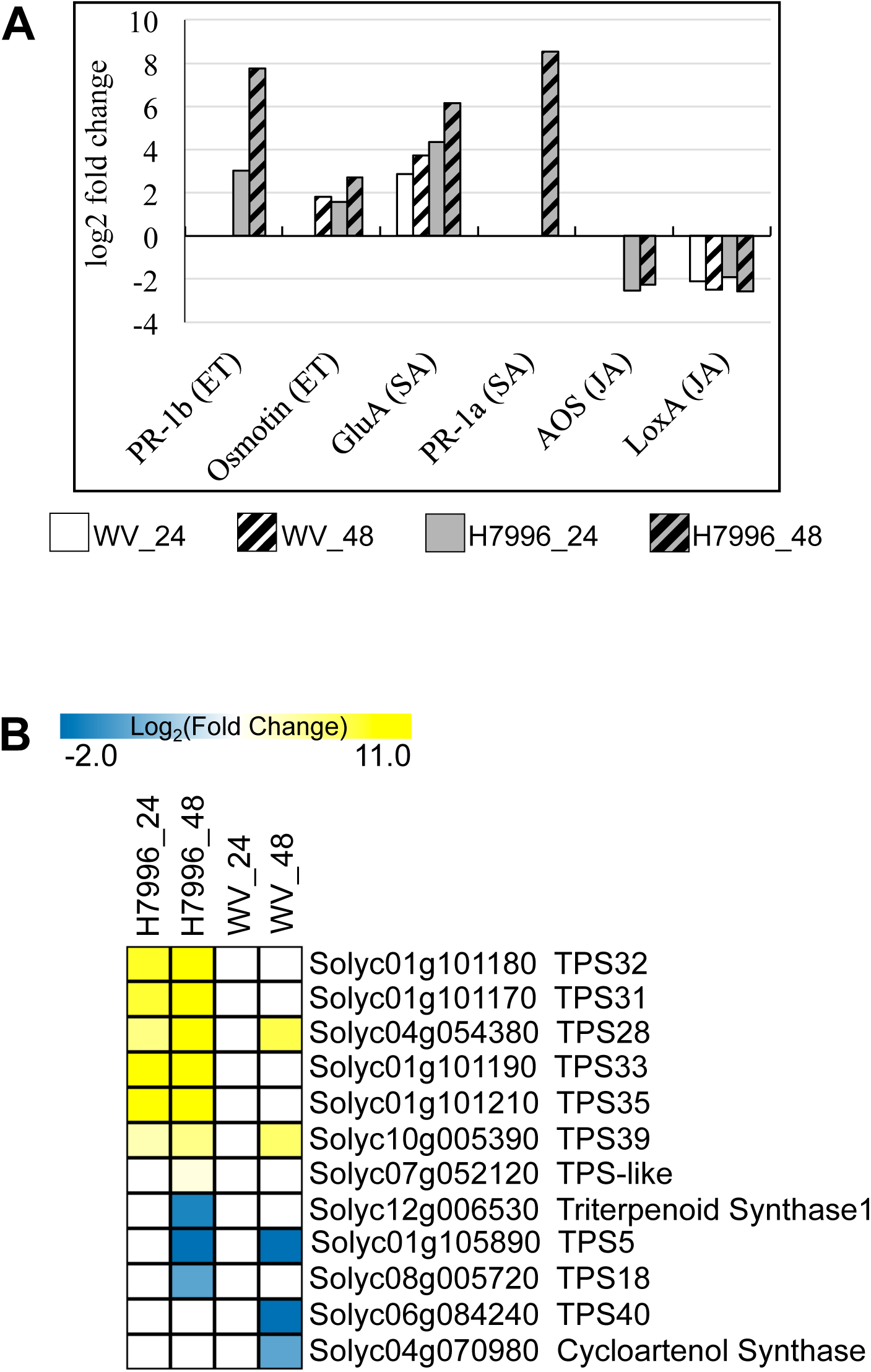
Defense responses are activated earlier and with higher fold changes in the root of resistant H7996. A) log fold changes in RNA-seq data of marker genes for classic defense hormones, B) Heat map showing log fold changes of genes in the ‘terpenoid’ bin in MapMan software (Thimm et al. 2004). More terpene synthase (TPS) genes are activated in roots of resistant plants and at an earlier time point.

Consistent with JA – SA antagonism (Robert-Seilaniantz et al. 2011; Derksen et al. 2013), and our GO analysis above, marker genes for JA defense responses were repressed in both resistant H7996 and susceptible WV, but showed greater fold change repression in roots of the resistant H7996 plants. *ALLENE OXIDE SYNTHASE* (*AOS*) and *LIPOXYGENASE* (*LoxA*) were both downregulated in resistant H7996 after both time points, *LoxA* was also downregulated in WV (Fig. 5a). This corresponded to the GO enrichment analysis that showed that regulation of JA mediated signaling was overrepresented in downregulated genes only for resistant H7996 (Fig. 4). Together, these results reveal activation of SA- and ET- dependent defense pathways earlier in roots of the resistant plant H7996, as well as an earlier deactivation of JA-dependent defense signaling in resistant H7996.

In addition to these classic defense pathways, we observed strong upregulation of terpene synthases in resistant tomato roots (Fig. 5b). Terpenoids are a large class of compounds composed of five carbon isoprene units, and are building blocks of some plant hormones and of specialized secondary metabolites (Falara et al. 2011). Tomato has 44 terpene synthase (TPS) genes, of which 29 are functional and are divided into 5 clades (Falara et al. 2011). In roots of resistant plants, five TPS genes in the alpha clade, which encode sesquiterpene synthases (TPS28, 31, 32, 33, 35), a TPS-like gene, and a linadool/nerolidol synthase (TPS39) are strongly upregulated at 24 hpi and 48 hpi (Fig. 5b). In contrast, only one sesquiterpene synthase, TPS28, and the linadool/nerolidol TPS39 are upregulated in susceptible roots at 48 hpi (Fig. 5b). Terpenoids act as antimicrobial or anti-insect compounds, and the strong upregulation observed in roots of resistant plants may contribute to resistance.

### Roots of susceptible tomato plants downregulate genes required for organ growth at 48 hpi

To have a better understanding of the response within roots of each genotype, we focused on genes that were exclusively responsive within each time point in each genotype (i.e. genes that were activated or repressed only within H7996 or WV at each time point, boxed numbers in Fig. 2b and c). All nine GO terms that overlapped among exclusive genes in WV and H7996 were related to defense and detoxification (Supplementary Fig. S2). Consistent with earlier and larger fold change defense responses in the resistant H7996 root, all but one of these categories were found both in genes upregulated in the resistant H7996 root at 24 hpi and genes downregulated in the susceptible WV root at 24 hpi (Supplementary Fig. S2).

Analysis of the 808 genes exclusively downregulated at 48 hpi in susceptible WV roots revealed several GO categories with known roles in root growth. These included GO categories ‘DNA replication’ (GO: 0006260; P = 8.7 × 10^-7^) (Ni et al. 2009; Jia et al. 2016), DNA packaging (GO:0006323; P = 4.4 × 10 ^-10^), chromatin assembly (GO:0031497, P = 9.7 × 10^-11^) (Shen and Xu 2009; Aichinger et al. 2011; Sang et al. 2012), and translation (GO: 0006412; P = 3.7 × 10^-31^) (Wieckowski and Schiefelbein 2012) (Fig. 6). Genes repressed in these categories included DNA replication helicases *MCM3* (*Solyc02g070780*), *MCM4* (*Solyc01g110130*), *MCM5* (*Solyc07g005020*) and *MCM7* (*Solyc01g079500*), ribosomal proteins and histones. In Arabidopsis, *MCM2* is involved in DNA replication and is important for root meristem maintenance (Ni et al. 2009), and mutations in a DNA helicase/nuclease result in very short roots (Jia et al. 2016). Further, mutation of AtMDN1, an AAA-ATPase that is a component of the pre-60S ribosome, results in several developmental defects including a shorter root (Li et al. 2016). Histone modifications have also been shown to be critical for proper root growth and development (reviewed in Takatsuka and Umeda 2015). None of these GO categories were identified within differentially expressed genes in the resistant H7996 root (Fig. 6).

**Fig. 6:**
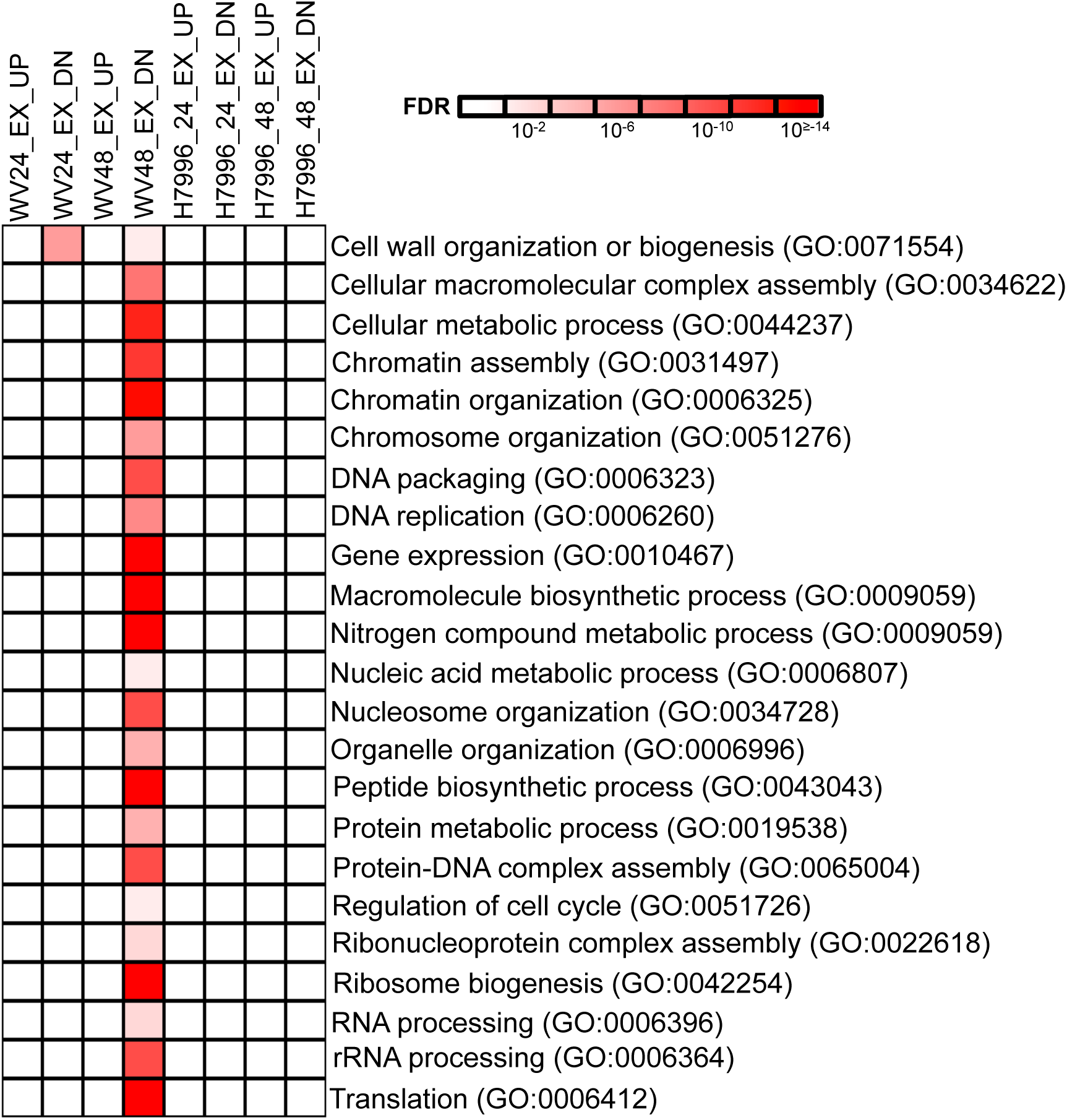
Roots of susceptible plants strongly repress pathways required for organ growth at 48 hpi. Heatmap of selected overrepresented GO categories (corrected P < 0.05) in up- and downregulated genes in roots of susceptible WV at 24 and 48 hpi. All GO categories in Supplementary Table 5. No overrepresented categories were observed in WV24_EX_UP.

These data suggested that roots of susceptible plants slow growth after infection. To test this, we quantified root growth of H7996 and WV at 10 dpi. Plants were removed from pots, and the root systems were gently washed with water to remove soil. Cleaned roots were scanned and surface area quantified using a WinRHIZO root scanning and quantification system (Arsenault et al. 1995). We find that roots of WV have significantly decreased surface area after inoculation compared to mock-inoculated controls (Fig. 7). In contrast, *R. solanacearum* inoculated roots of resistant H7996 have no difference in surface area compared to mock-inoculated resistant roots (Fig. 7). The differential root growth response to *R. solanacearum* between resistant and susceptible accessions is consistent with the transcriptional changes that we observed.

**Fig. 7:**
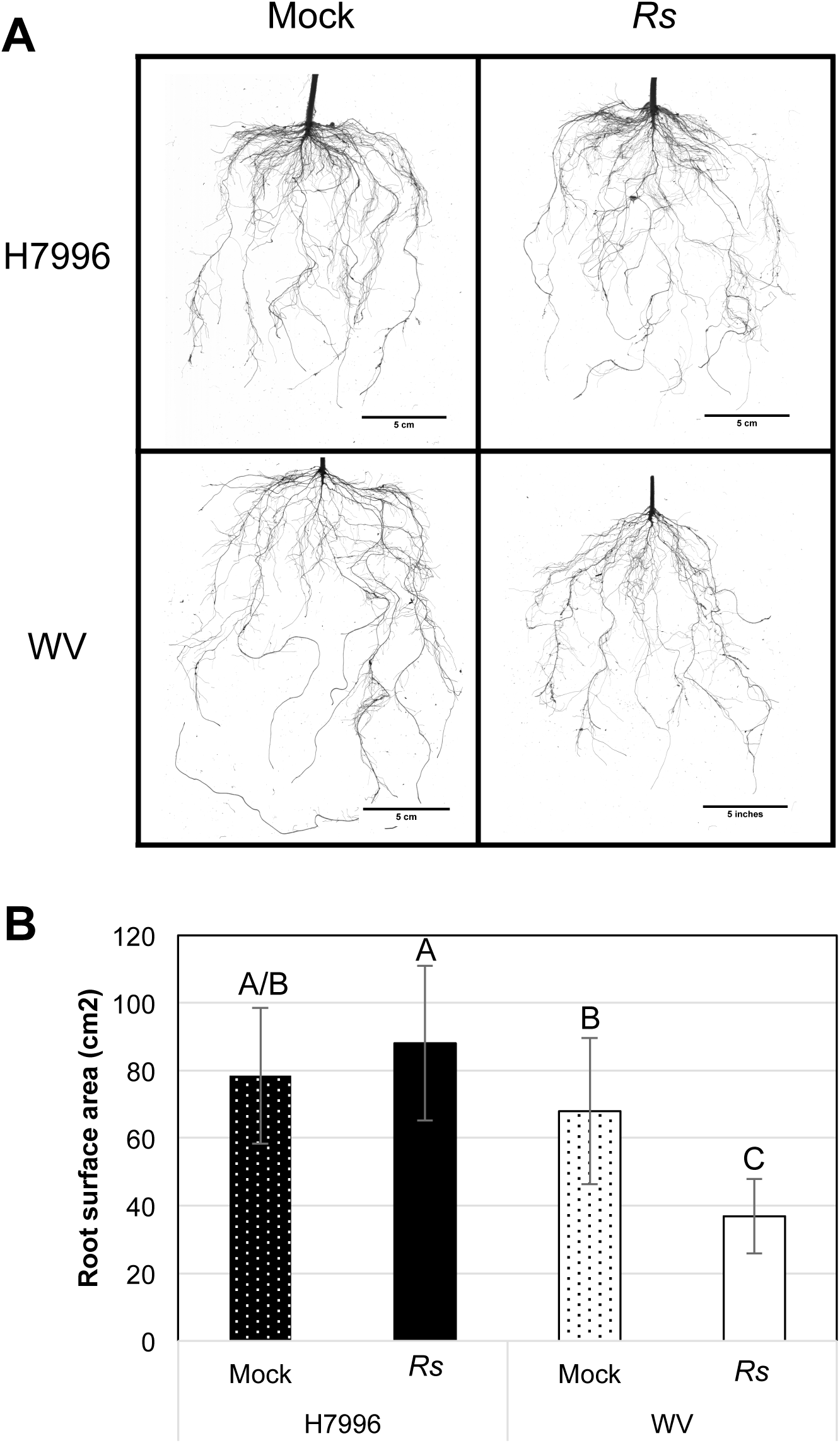
Root architecture of resistant H7996 and susceptible WV at 10 dpi. A) *R. solanacearum* (*Rs*) and mock-inoculated roots at 10 dpi imaged with a flatbed scanner. Representative images from three independent experiments, each with at least five roots per genotype and treatment, are shown, B) Quantification of whole root surface area using the WinRHIZO software image analysis system (Arsenault et al. 1995). Letters indicate significant differences (P < 0.05) with a two-way ANOVA and Tukey’s HSD test.

Consistent with the hypothesis that the susceptible WV root, responds to *R. solanacearum* with growth suppression, far fewer GO categories were overrepresented in the set of exclusively upregulated genes in WV roots at 48 hpi (Supplementary Table S4). Three GO categories were identified among the 594 number of genes exclusively upregulated in WV, compared to 72 categories identified among the 808 downregulated genes. Among the three GO categories overrepresented in the exclusively upregulated genes in WV at 48 hpi was ‘defense response’ (GO: 0006952; P = 1.01 × 10^-4^) (Supplementary Table S5). Together these results show that although roots of the susceptible WV plant do eventually activate defense responses, they are also initiating processes that limit root growth.

### Auxin response pathways are altered in roots of resistant plants

GO analysis of genes that were exclusively expressed in roots of the resistant variety H7996 at each time point revealed that the categories ‘auxin-activated signaling pathway’ (GO:0009734; P = 4.3 × 10^-2^) and ‘cellular response to auxin stimulus’ (GO: 0071365, P = 4.3 × 10^-2^) were overrepresented in genes exclusively downregulated in the resistant H7996 at 48 hpi (Fig. 8).

**Fig. 8:**
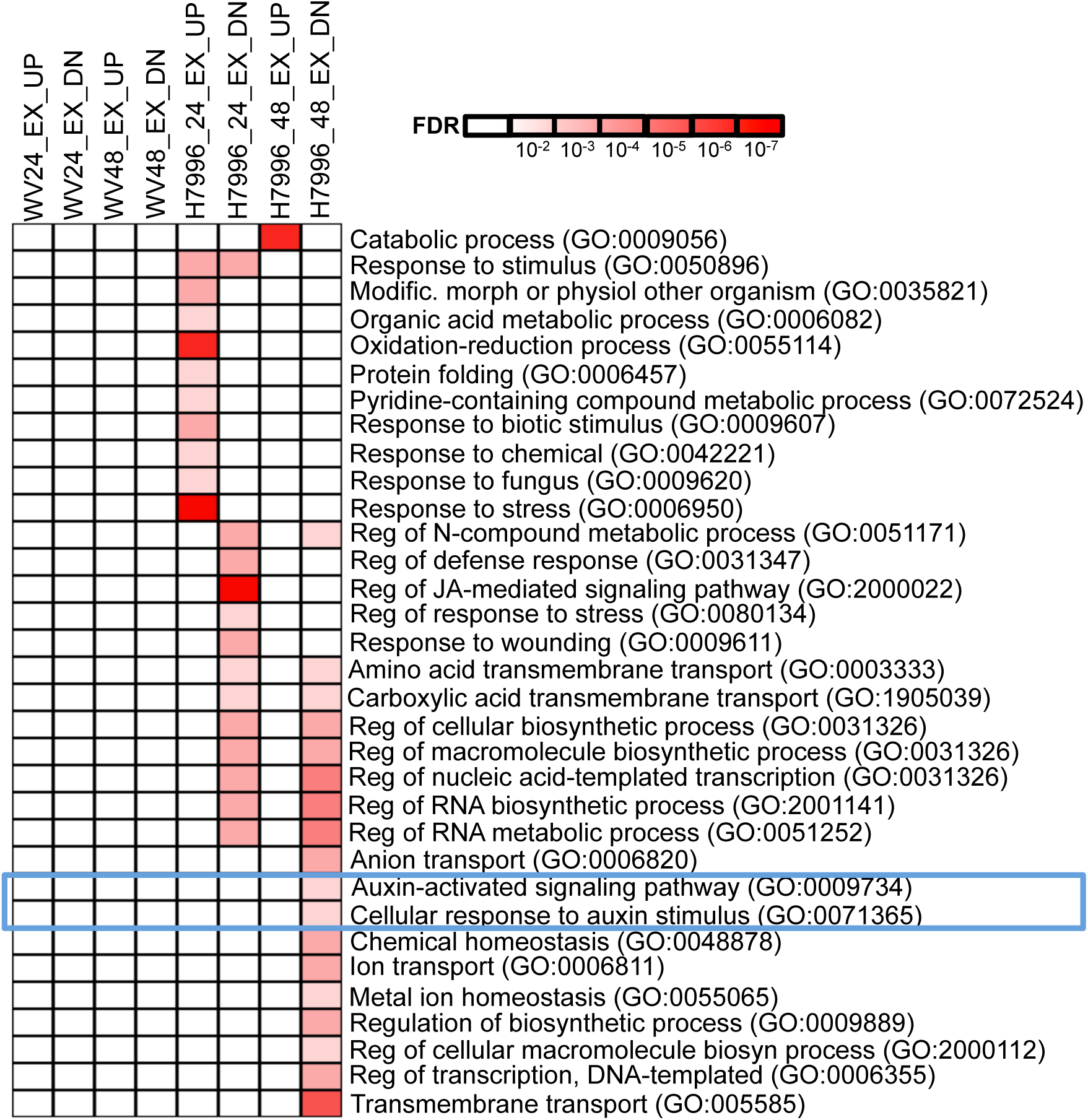
Auxin-related and lateral root development genes are differentially expressed in the resistant root at 48 hpi. Selected GO categories overrepresented among genes exclusively differentially expressed in H7996 at each of the time points shown. The blue box highlights auxin-related GO categories. The nine categories that overlapped between H7996 and WV are shown in Supplementary Fig. 2 and are not shown here.

Examination of the eight genes within these categories identified three genes encoding transcription factors known as *AUXIN RESPONSE FACTORs* (*ARF*s), which have both positive and negative roles in auxin signaling. These included two *S. lycopersicum* orthologs (*Solyc12g042070* and *Solyc03g118290*) of Arabidopsis *ARF2*, and the *S. lycopersicum* ortholog of Arabidopsis *ARF4* (*Solyc11g069190*). Of the other five genes within the ‘auxin response’ GO category, one encoded a PIN auxin transporter (*Solyc10g080880*), three were AUXIN/INDOLE-3-ACETIC-ACID (AUX/IAA) transcription factors (*Solyc06g008590*, *Solyc06g008580*, *Solyc01g097290*), and another encoded an uncharacterized gene (*Solyc02g036370*) related to the REVEILLE1 transcription factor in Arabidopsis.

### The tomato auxin transport mutant diageotropica (dgt) is resistant to R. solanacearum

One of the genes within the auxin response GO category above was *Solyc10g080880,* which encodes a PIN auxin efflux transporter known as SISTER OF PIN1b (SlSoPIN1b). PIN proteins are the primary auxin efflux transporters in plants and are responsible for polar auxin transport (Krecek et al. 2009; Adamowski and Friml 2015). In Arabidopsis, mutations in several auxin transporters, including *PIN2*, lead to decreased disease symptoms caused by *Fusarium oxysporum* (Kidd et al. 2011). We hypothesized that tomato genes required for polar auxin transport function in resistance to *R. solanacearum*. To test this, we examined resistance of the tomato mutant *diageotropica* (*dgt*) to *R. solanacearum*. *DGT* encodes a cyclophilin that negatively regulates PIN auxin efflux transporters in tomato (Ivanchenko et al. 2015). Mutations in *DGT* lead to altered auxin transport and changes to the transcription and/or protein localization of PINs (Ivanchenko et al. 2015). Root inoculation of the *dgt1-1* mutant and its susceptible wild type parent, Ailsa Craig (AC), showed that *dgt1-1* was highly resistant to *R. solanacearum* compared to the wild type parent (Fig. 9). Three independent biological replicates revealed that mutant plants had consistently less than 10% wilting at 12 dpi. In contrast, the wild type parent had almost 80% wilting at the same time point.

**Fig. 9:**
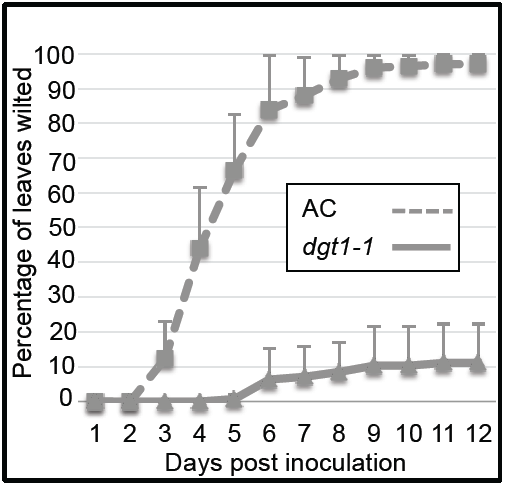
The *dgt1-1* mutant shows enhanced resistance to *R. solanacearum* compared to its wild type control AC with root soaking inoculation. Wilting was scored daily based on the percentage of leaves wilted per plant. Each point represents the average of three independent experiments, each with 8 - 9 plants per genotype. Area Under the Disease Progress Curve (AUDPC) for AC = 725.2 ± 85.2 and for *dgt1-1* = 60 ± 64.2 (P < 0.001 with a two-tailed t-test). Error bars indicate standard deviation.

### The increased resistance of *dgt1-1* is not due solely to alterations in root architecture

The *dgt1-1* mutant has been previously described as lacking lateral roots (Muday et al. 1995; Oh et al. 2006; Ivanchenko et al. 2015). Because *R. solanacearum* enters the root system in part through wounds created as lateral roots emerge from the primary root, we questioned whether the decreased colonization of *R. solancearum* in *dgt1-1* was due to deficiencies in lateral root emergence. Previous work showing a lack of lateral roots in *dgt1-1* used plants grown in agar (Ivanchenko et al. 2015). However, examination of root systems of *dgt1-1* grown in potting mix revealed that the mutant does produce lateral roots in these conditions (Fig. 10B, arrows), although roots of *dgt1-1* were still significantly smaller compared to the wild-type parent AC (Fig. 10).

**Fig. 10:**
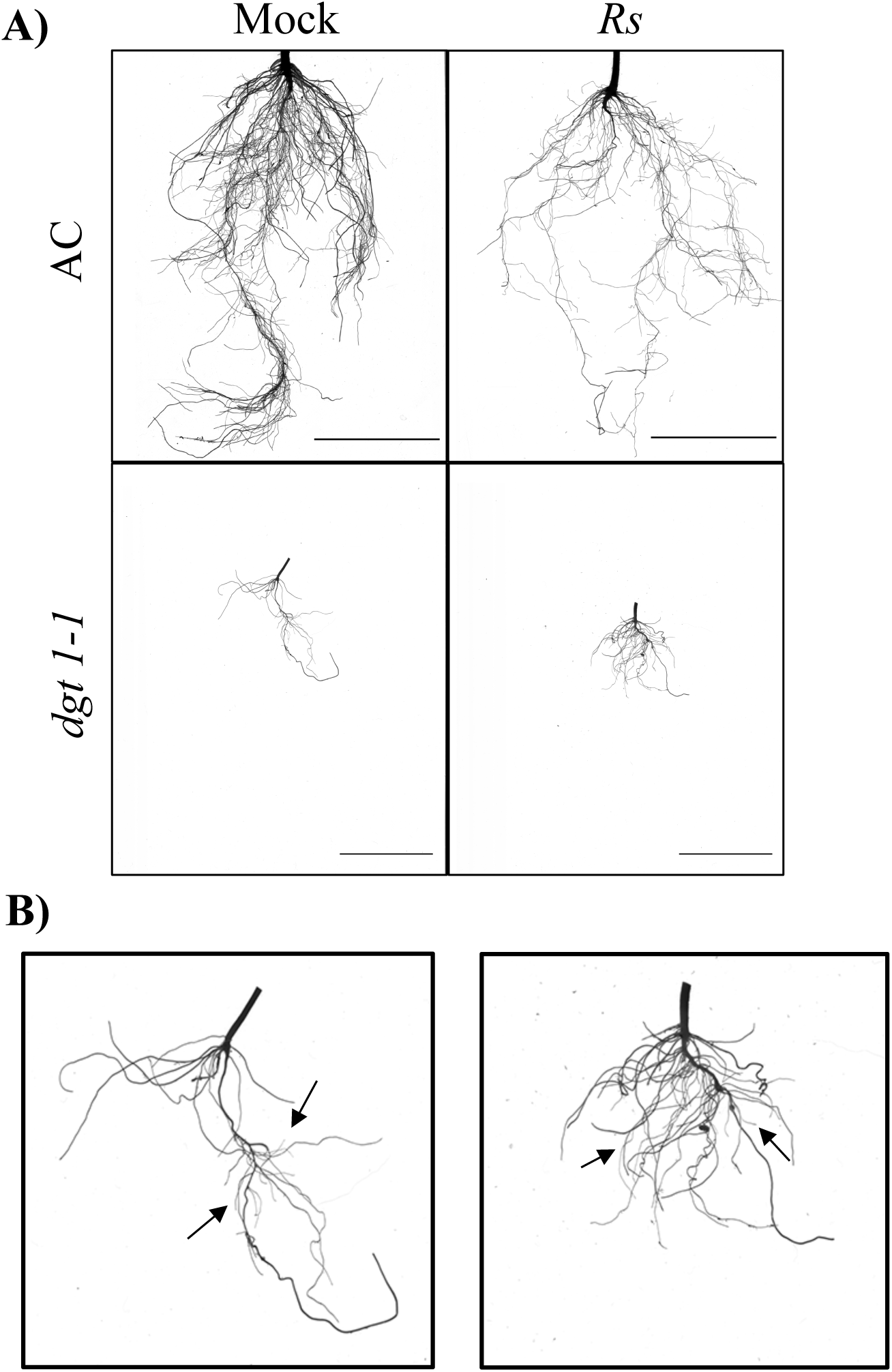
Root architecture of susceptible AC and resistant *dgt1-1* at 6 dpi grown in potting mix and soil-soak inoculated with either water (mock) or *R. solancearum* strain K60 (*Rs*). A) Plants were grown in potting mix and roots imaged with a flatbed scanner, B) Close-up images of *dgt1-1* plants in (A). Arrows point to examples of lateral roots. Images are representative of those from two independent biological replicates with six plants per replicate per treatment and genotype. Scale bars = 5 cm.

To examine whether the altered root structure was the underlying basis for the increased resistance, we used petiole inoculation of *R. solanacearum* in the *dgt1-1* and AC mutant. This method bypasses the root system by directly injecting bacteria into the petiole vasculature (Tans-Kersten et al. 2001; Dalsing and Allen 2014). If decreased lateral root emergence in the *dgt1-1* mutant were the primary reason for resistance, we would expect that the *dgt1-1* mutant would show an increased susceptibility using this method. Using petiole inoculation, the *dgt1-1* mutant did not wilt by 12 dpi, compared to approximately 90% wilting in the wild type AC control (Fig. 11). Together, these results suggest that the enhanced resistance to *R. solanacearum* in the *dgt1-1* mutant is due to modulation of auxin transport.

**Fig. 11:**
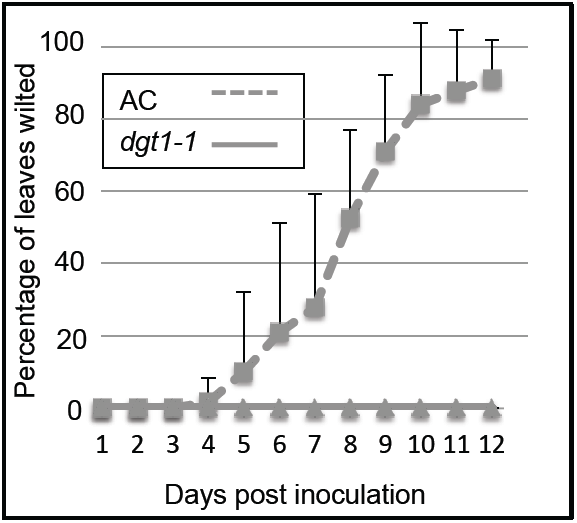
The *dgt1-1* mutant shows enhanced resistance to *R. solanacearum* compared to its wild type susceptible parent AC with petiole inoculation. Wilting was scored daily based on the percentage of leaves wilted per plant. The experiment was repeated three times with 3 – 9 plants of each genotype per experiment. The average of three experiments is shown. The average Area Under the Disease Progress Curve (AUDPC) for AC = 401.6 ± 154.8; average AUDPC for *dgt1-1* = 0 ± 0 (P < 0.01; two-tailed t-test with unequal variance). Error bars represent standard deviation.

## DISCUSSION

In this manuscript we show that infection with the soilborne pathogen *R. solanacearum* leads to a strong defense response in tomato roots that includes alteration of auxin pathways. Analysis of a tomato mutant with defective auxin transport confirmed a role for auxin pathways in resistance. Susceptible tomato roots are stunted at 6 dpi, and consistent with this, we find significant suppression of genes required for growth and cellular homeostasis at 24 and 48 hpi. Additionally, roots of the susceptible variety are slower to activate defense responses, and their defense responses are lower in magnitude compared to resistant roots.

Genome-wide transcriptional responses to *R. solanacearum* in tomato have been previously examined primarily in aboveground regions of the plant. (Ghareeb et al. 2011; Ishihara et al. 2012; Kiirika et al. 2013). Ishihara *et al*. 2012 used tomato microarrays to examine gene expression changes 24 hpi with *R. solanacearum* strain 8107S in stems and leaves of susceptible tomato cultivar Ponderosa and resistant LS-89. They did not identify any changes in gene expression at 24 hpi in the susceptible cultivar, and only 143 genes were differentially expressed in leaves of the resistant cultivar compared to the mock-inoculated controls. Differences in our results can be explained in part by the region of the plant sampled (aboveground vs. belowground), inoculation method, or the result of differences in the gene expression profiling method used in each study (microarray vs. RNA-seq). Despite these differences, several of the genes upregulated in resistant tomato stems were found in similar pathways as those we identified in roots of resistant H7996, including *PR* genes. In line with the idea of some overlap in defense responses between below and aboveground regions to *R. solanacearum*, defense marker gene expression in aboveground regions of resistant tomato plants also occurred earlier and more strongly in resistant H7996 compared to susceptible variety Bonnie Best (Milling et al. 2011). Together, these data suggest that root defense responses partially overlap with those in the shoot, but also have unique responses to pathogen attack.

We observed a strong upregulation of terpene synthase genes specifically in roots of resistant plants. Analysis of ginger leaves after rhizome infection with *R. solanacearum* revealed a similar upregulation of terpene synthases in resistant plants (Prasath et al. 2014). A previous report (Lin et al. 2014) used virus-induced gene silencing in resistant H7996 to knock down expression of four TPS genes (*TPS31*, *TPS32*, *TPS33*, and *TPS35*) that were highly upregulated in our dataset. They found that more silenced plants were colonized by *R. solanacearum* in the stem, suggesting that TPS silenced lines had decreased tolerance to *R. solanacearum*. These data suggest that upregulation of TPS genes may contribute to resistance in tomato and ginger. However, this does not appear to be a mechanism used in all crops, as in peanut, terpenoid synthase genes were downregulated at 12 hpi after infection in both resistant and susceptible genotypes (Chen et al. 2014). Indeed, resistance in peanut may operate through different mechanisms than in tomato, as evidenced in the root of a resistant peanut genotype, in which many NBS-LRR type resistance genes and genes encoding proteins with a LRR-LRK motif were mainly downregulated (Chen et al. 2014).

Our data show both commonalities and differences in resistance between tomato variety H7996 and wild potato species *S. commersonii* (Zuluaga et al. 2015). In resistant roots of both species, more genes with roles in biotic stress were upregulated than downregulated. However, in contrast to our results, which found overrepresentation of the JA pathway in downregulated genes of resistant roots, no genes in the JA pathway were downregulated in roots of resistant potato plants (Zuluaga et al. 2015). Additionally, in resistant wild potato roots, genes in the auxin pathway were upregulated and none were repressed (Zuluaga et al. 2015), while we observed overrepresentation of auxin pathways in downregulated genes in resistant tomato roots. These differences could be the result of differences in species, or to time of inoculation, as we sampled our plants at an earlier time point (24 and 48 hpi compared to 3 – 4 days).

Suppression of auxin biosynthesis, responses and signaling has been associated with plant resistance to biotrophic or hemi-biotrophic pathogens in multiple pathosystems (reviewed in (Fu and Wang 2011; Ludwig-Muller 2015)). In Arabidopsis, mutations in several auxin transporters, including *PIN2* and *AUX1*, reduce disease severity caused by the pathogenic fungus *Fusarium oxysporum* (Kidd et al. 2011). The *walls are thin* (*wat1*) mutant of Arabidopsis is resistant to *R. solanacearum*, has decreased auxin content in roots, suppressed indole metabolism, and decreased tryptophan in roots at 4 dpi (Denance et al. 2012). *WAT1* encodes a vacuolar auxin transporter (Ranocha et al. 2013) and appears to modulate both cellular auxin levels within the vascular tissues as well as whole organ levels of auxin in the root and stem. Intriguingly, *wat1* is resistant to multiple pathogens that, like *R. solanacearum*, colonize the vasculature, but not to non-vascular pathogens such as *Pseudomonas syringae* pv. *tomato* (Denance et al. 2012). Resistance to *R. solanacearum* was dependent on SA, because *wat1 NahG* plants showed comparable levels of disease to wild type Arabidopsis. The *wat1* mutant was first identified due to a defect in secondary cell wall biosynthesis (Ranocha et al. 2010). Mutations in genes required for secondary cell wall formation including *CELLULOSE SYNTHASE4* (*CESA4*)/*IRREGULAR XYLEM5* (*IRX5*), *CESA7*/*IRX3*, and *CESA8*/*IRX1*, also lead to enhanced resistance to *R. solanacearum* in Arabidopsis (Hernandez-Blanco et al. 2007). However, in these mutants, resistance is independent of the SA pathway, but dependent on ABA responses (Hernandez-Blanco et al. 2007).

Here we showed that genes in auxin pathways, including *SlSoPIN1b*, a homolog of the PIN1 auxin transporter, are overrepresented in exclusively downregulated genes in resistant tomato roots after *R. solanacearum* infection. We find that a tomato mutant with altered auxin transport is resistant to *R. solanacearum*. Mutations in tomato *DGT* lead to changes in polar auxin transport that result in abnormal auxin distribution along the root (Ivanchenko et al. 2006). Polar auxin transport is crucial for plant development and is mediated by PIN auxin transporters (reviewed in (Krecek et al. 2009; Adamowski and Friml 2015). Roots are composed of multiple cell types and tissues that differ in auxin levels (Petersson et al. 2009). In Arabidopsis, most PIN transporters localize to the plasma membrane on specific faces of the cell, and their localization varies depending on root cell type (Blilou et al. 2005). The tomato DGT protein regulates levels and localization of PIN1 and PIN2 transporters in the root (Ivanchenko et al. 2015). In wild type tomato roots, PIN1 localizes to the rootward face of cells in the root stele (Ivanchenko et al. 2015). The *dgt* mutation leads to decreased PIN1 protein in the stele of root tips. In addition, expression of *PIN2* is significantly decreased in root tips of the *dgt* mutant and the PIN2 protein localization is altered (Ivanchenko et al. 2015). Although auxin levels in whole roots of the *dgt* mutant are greater than those in wild-type plants (Ivanchenko et al. 2006), auxin responses and signaling in the root vasculature are decreased (Ivanchenko et al. 2015) due to the altered localization of PIN1 and PIN2. How mutations in *DGT* lead to resistance is not entirely clear. One possibility is that resistance is due to antagonism between auxin and SA. Alternatively, like *wat1* and other Arabidopsis mutants, *dgt* may have altered secondary cell wall structure that enhances resistance, or may be altered in another auxin-related process that results in enhanced resistance.

Understanding mechanisms of root-mediated resistance is an important step in developing crops with resistance to soilborne pathogens. Like many other bacterial pathogens, *R. solanacearum* produces auxin (Valls et al. 2006). Whether resistant plants downregulate auxin pathways to overcome pathogen auxin production, and whether the alteration of auxin transport is a general feature of root-mediated resistance are intriguing questions whose answers may lead to new insights into enhancing crop resistance.

## MATERIALS AND METHODS

### Plant growth and *R. solanacearum* K60 inoculation

Resistant tomato (*Solanum lycopersicum* L.) accession Hawaii7996 (H7996), susceptible *pimpinellifolium* West Virginia 700 (WV), *digeotropica* (*dgt1-1; S. lycopersicum*), and Ailsa Craig (AC; *S. lycopersicum*) were grown in Propagation Mix (Sun Gro Horticulture) in square pots containing 25-27g of soil and grown under 16:8 h light, 28° - 30°C in a growth chamber. The *dgt1-1* mutant has been previously reported (Oh et al. 2006), and we confirmed that the mutation was present by sequencing the gene. Growth and inoculation of *R. solanacearum* was as described in (Caldwell et al. 2017). Briefly, *R. solanacearum* strain K60 (phylotype IIA, sequevar 7) was recovered from a glycerol stock and grown for two days on Casamino Peptone Agar (CPG) containing 1% triphenyltetrazolium chloride (TZC) at 28°C. Bacteria were harvested with sterile water and resuspended to 1.0 × 10^8^ CFU/ml. At the three-leaf stage (approximately 14 – 17 days after planting), tomato plants were root-inoculated by gently lifting plants from their growth containers, and then soaking in either inoculum or water to the root-shoot soil line (approximately 40 ml per plant)(as in Caldwell et al. 2017). After soaking for 5 min, seedlings were transferred back to their growth containers and placed back into a growth chamber with the conditions above. Dilution plating was used to confirm the concentration of inoculum after each set of inoculations.

For *dgt* and AC resistance tests, wilting was rated daily and scored as the percentage of leaves per plant wilted. For each of soil-soak and petiole inoculation, average wilting with standard error are shown for three independent experiments. For soil soak inoculation, each independent experiment had 8 - 9 plants per genotype, and for petiole inoculation, each experiment had 3 – 9 plants per genotype. The Area Under the Disease Progress Curve (AUDPC) was calculated according to (Madden et al. 2007) with percent leaf wilting used as the disease measure.

### Plant colonization assays

Individual plants from both mock and *R. solanacearum* inoculations were removed from pots, and the soil was gently washed off in a tray of sterilized distilled water. Roots of each plant were transferred into a 50 ml Falcon tube containing 45 ml of sterilized distilled water, and further cleaned to remove residual soil by shaking the Falcon tube for 1 minute. This wash was repeated 5 times. Water from cleaned roots was removed with a dried paper towel and roots were weighed. Washed, cleaned roots were surface sterilized by dipping in 100% ethanol for 30 seconds, and then flamed quickly to remove residual ethanol. Each surface sterilized root was ground in 1 ml ddH_2_O with a mortar and pestle, the lysate was centrifuged briefly, and the supernatant was used to determine *R. solanacearum* K60 titer with serial dilutions in ddH_2_O. 100 μl of diluent was plated on CPG plates containing 1% TZC and incubated at 28°C for 48 hours. Colonies were counted and *R. solanacearum* K60 titer was determined as CFU/g of tissue. Colonization assays were performed in three independent experiments with three plants per genotype and time point per experiment. Data did not meet the assumption of normality and the Mann Whitney Wilcoxon test was performed in RStudio version 0.99.484.

### Total RNA extraction and RNA-seq sample preparation

Whole roots from 10 plants of each genotype (H7996 and WV) were harvested at each time point (0 hour mock-inoculation, 24 hpi, and 48 hpi). Roots from these 10 plants were pooled for each genotype at each time point in each replicate. Three independent replicates were performed. Samples were ground into a powder using a mortar and pestle under liquid nitrogen. 100 mg of ground root tissue from each sample was used for total RNA extraction using Trizol following the manufacturer’s instructions (Invitrogen, CA). 50 μg of extracted total RNA was subjected to RNAse-free DNase (Omega, GA) treatment. DNAse treated total RNA was further cleaned using a Nortek column following the manufacturer’s instructions (Norgen BioTek Corp., Canada). Two μg from each of 18 samples (three time points x two genotypes x three replicates) were submitted to the Purdue Genomic Center for RNA-seq on the Illumina HiSeq 2500. RNA quality was determined using an Agilent Nanochip (Agilent, CA) and all samples had a RIN score of at least 7.8. Stranded mRNA libraries were constructed at the Purdue Genomics Facility using Illumina’s TruSeq Stranded mRNA Sample Preparation kit (Revision E, Oct 2013) according to the manufacturer’s instructions.

### RNA-seq data analysis

Illumina paired-end 100 bp RNA sequencing was performed on all samples. A total of 967,730,337 reads were generated after quality filtering and mapping (Supplementary Table S1). Reads for each of the 18 samples were aligned by the Purdue Genomics Facility to the ITAG2.4 *S. lycopersicum* reference genome using Tophat2 version 2.0.14 (Trapnell et al. 2009). Library type was set to strand-specific (first strand), mate inner distribution to 300, and mate standard deviation to 150. Gene expression was measured as the total number of reads for each sample that uniquely mapped to the reference, binned by gene. Each sample averaged about 54 million high quality, uniquely aligned reads (Supplementary Table S1). After filtering for low counts such that at least 3 of the 18 samples had at least 3 counts per million (CPM) per row, a total of 20,641 genes remained for differential expression analysis. Differential gene expression analysis was performed using the edgeR package (Robinson et al. 2010) in Bioconductor version 3.3. The edgeR function calcNormFactors was used for library normalization with the default edgeR trimmed mean of M-values (TMM) method. Normalized library sizes are listed in Supplementary Table S2. Differentially expressed genes were identified using the glm (General Linear Model) pipeline in edgeR according to the edgeR documentation. The design matrix was created with coefficients for the expression level of each group. A group consisted of genotype and time point (H7996_0 hour = group 1, H7996_24 hour = group 2, etc). Common and tagwise dispersions were estimated with the function estimateDisp function. Multidimensional scaling (MDS) analysis revealed no batch effect of different replicates (Supplementary Figure S3).

Pairwise comparisons were performed between mock 0 hour and 24 hpi, and between mock 0 hour and 48 hpi within each of H7996 and WV using the contrast argument in the glmLRT function. Differential expression was determined using the Benjamini-Hochberg false discovery rate (FDR) multiple testing correction (Benjamini and Hochberg 1995) with an adjusted P-value of 0.05 and a log2 fold change > |0.585| (corresponds to a fold change of > |1.5|). Venn diagrams were generated using VENNY 2.1 (Oliveros 2007-2015). Gene ontology (GO) analysis was performed using the PANTHER GO analysis tool (http://www.pantherdb.org/) (Huaiyu et al. 2016). GO terms are derived from annotations of the sequenced *S. lycopersicum* genome, Heinz1706 (Tomato Genome Sequencing Consortium 2012). All GO categories shown are for ‘biological process’. Heat maps, including those for GO figures were visualized with Multiple Experiment Viewer from TM4 (Saeed et al. 2003; Saeed et al. 2006).

### cDNA synthesis and qRT-PCR

Total RNA extraction was performed as above from root tissue used in the RNA-seq analysis. cDNA synthesis and qRT-PCR was performed as in (Kim et al. 2017). Two biological replicates were used for validation. Briefly, cDNA was reverse-transcribed from 1 μg RNA using the NEB AMV first strand cDNA synthesis kit as per manufacturer’s instructions. Quantitative RT-PCR was performed with 1μl of cDNA on a Roche Light Cycler (Roche, CA) with the following amplification protocol: 50°C for 2 min and 95°C for 2 min followed by 40 cycles of 95°C for 15 sec and 60°C for 1 min. PCR efficiency of the primers ranged from 95 % to 105 %. *ACTIN* (*Solyc11g005330*) was used as the gene for normalization. *Solyc11g005330* was not differentially expressed in either H7996 or WV at either time point (Supplemental Table 4). The ΔΔCt method (Livak and Schmittgen 2001) was used to calculate fold changes relative to the internal control and the mock-inoculated control plant. Primer sequences are listed in Supplementary Table S5.

### Root architecture measurements

Roots were harvested from mock and *R. solanacearum* -inoculated plants at 6 dpi (AC and *dgt1-1*) or 10 dpi (WV and H7996). Whole root systems were washed gently in water and scanned with a calibrated color optical scanner from Regent Instruments, Inc (Quebec, Canada) and measured using software in the WinRHIZO V. 2016a system (Regent Instruments Inc, Quebec, Canada) (Arsenault et al. 1995). Data were analyzed with a two-way ANOVA followed by post-hoc Tukey’s honest significant differences (HSD) test using RStudio version 0.99.484. No transformations were necessary to meet the homogeneity of variance and normality assumptions. Two independent biological replicates with at least six plants per treatment and genotype were performed for AC and *dgt1-1*. Three independent biological replicates with at least 5 roots per treatment and genotype were performed for WV and H7996. Representative images are shown.

## ACKNOWLEDGEMENTS

We thank Maria Ivanchenko for the *dgt* mutant, Erin Sparks and members of the Iyer-Pascuzzi lab for comments on the manuscript, and Pete Pascuzzi for helpful comments on RNA-seq analysis. This work was funded by a grant from the Foundation for Food and Agriculture Research, start-up funds from Purdue University, and Hatch funds to AIP, and a NSF Graduate Research Fellowship DGE-1333468 (to E.F).

## SEQUENCE DATA

All RNA-seq data in this study has been submitted to the Short Read Archive (SRA) at NCBI under project number SRP078159. The SRA does not provide pre-release access to sequence data, but a reviewer link to the metadata is here: ftp://ftp-trace.ncbi.nlm.nih.gov/sra/review/SRP078159_20170811_115714_3d522deaf85577451c01974654b36ad3

## SUPPORTING INFORMATION LEGENDS

Supplementary Fig. S1: Comparison of qRT-PCR and RNA-seq data at 24 and 48 hpi after inoculation in H7996 and WV. Two biological replicates were used for the qRT-PCR analysis. Expression of genes is shown for both genotypes and time points only if the gene expression was significant (FC > 1.5, q < 0.05) in the RNA-seq data. Error bars show standard deviation for qRT-PCR data.

Supplementary Fig. S2: GO categories that are found in more than one ‘Exclusive gene’ list.

Supplementary Fig. S3: Multidimensional Scaling (MDS) plot of RNA-seq samples. To examine the tomato root response to *R. solanacearum* inoculation within resistant H7996 and susceptible WV, differentially expressed genes were identified from two comparisons within each genotype: 1) 24 hpi to 0 hour and 2) 48 hpi to 0 hour.

Supplementary Table S1: Mapping summary showing numbers of raw reads, quality and adaptor clipped reads, TopHat mapping percentage for each of the 18 RNA-seq samples, and specified options in TopHat.

Supplementary Table S2: Normalized library sizes Supplementary Table S3: Raw counts of RNA-seq data.

Supplementary Table S4: Mean CPMs, log Fold Changes, logCPMs, LR, P values and FDR values from edgeR.

Supplementary Table S5: Gene Ontology (GO) categories for biological process from PANTHER for each time point comparison.

Supplementary Table S6: Primers used for qRT-PCR analysis.

